# Helping can be driven by non-affective cues in rat

**DOI:** 10.1101/2022.07.01.498150

**Authors:** Y. Vieira Sugano, H.Z. Shan, N.M.R. Molasky, P. Mason

## Abstract

Helping another in distress can be motivated by either affective or cognitive empathy, with the latter commonly believed to be restricted to humans and possibly other apes. Here, we found evidence for rodent helping that occurs in the absence of affective cues. We employed a paradigm in which a free rat can open the door to a restrainer containing a trapped rat. When the trapped rat was treated with the anxiolytic midazolam, the helping behavior exhibited by the free rat was diminished but did not extinguish. Correspondingly, midazolam-treated trapped rats still released themselves when given the opportunity, albeit at longer latencies than controls, evidence that midazolam only partially reduced the distress experienced by trapped rats. To test whether helping could occur for a rat who exhibited no affect, trapped rats were immobilized by general anesthesia or heavy sedation. Surprisingly, rats opened the door to restrainers containing these immobilized rats and pulled the incapacitated rats out of the restrainer, pushing them away from the arena center. The same solicitous behavior was observed when an anesthetized rat was simply placed in the center of the arena, without being confined within a restrainer. We speculate that the cognitive dissonance of immobile rats, at odds with predictive expectations of rat behavior built up over a lifetime, motivated solicitous behavior including helping. To block affective behavioral displays without associated dissonant cues of immobility, metyrapone, a drug that selectively blocks corticosterone synthesis, was administered to trapped rats. Under such circumstances, little helping behavior occurred. In sum, rats may be motivated either by affect or by cognitive dissonance, the latter comprising a rudimentary form of cognitive empathy.

## Introduction

Rodent helping exhibited towards conspecifics in distress is thought to depend on the social communication of mental state, which has been termed empathy (Preston & de Waal, 2002; Meyza et al 2017). The dominant model holds that the affective state of a distressed rat is “caught” by a potential helper rat, a basic form of empathy that has been called emotional contagion. Both mice and rats exhibit behaviors indicative of affective states such as fear-associated freezing solely based upon exposure to demonstrative conspecifics, i.e. independent of any direct fear-inducing experience such as foot shock (Atsak et al 2011; Panskepp & Lahvis, 2011; Jeon et al 2010; Burkett et al 2016; Meyza et al 2017; Langford et al 2006). On the other hand, using cognitive communication that does not rely upon affect as a motivation for helping has been considered beyond the capacity of the rodent brain. This untested bias has left the contribution of non-affective cues to rodent prosocial behavior largely unexplored.

In the trapped rat helping paradigm, a free rat is placed in an arena with a rat trapped in a centrally located restrainer with a door that can be opened only from the outside (Ben-Ami Bartal et al 2011). Within a few days of daily testing, most free rats start to open the door consistently and at short (<5 min) latency. Door-opening frees the trapped rat and only develops in the presence of a trapped rat. Door-opening does not depend on social interaction between the free and liberated rats (Ben-Ami Bartal et al 2011, Sato et al 2015, Cox and Reichel 2020).

Rats treated with midazolam (mdz), a benzodiazepine anxiolytic, do not develop door-opening, an indication that helper rats are motivated by affect (Ben-Ami Bartal et al 2016). As the source of the free rat’s affective distress, the trapped rat’s distress is expected to be greater than, or at least equal to, that expressed by the free rat. However, in experiments conducted in parallel and reported here, mdz treatment of the trapped rat had a far milder effect on helping than does such treatment of the free rat. One possible reason for this finding is that the trapped rat’s affect remains high and is minimally affected by the mdz dose used. Indeed, in experiments with a reversed door reported here, trapped rats, even those treated with mdz, free themselves consistently and at very short latency.

A second non-exclusive possibility is that free rats are motivated by non-affective, termed cognitive, cues to open the door. To test this, we blocked all affective expression by immobilizing trapped rats with either general anesthesia or a sedating dose of mdz. We found that the untreated free rats opened the restrainer door, dragged out the immobile trapped rats, and pushed them out of the arena center; such solicitous behaviors occurred even when immobile rats were not trapped. To determine whether the situational cue of a (non-distressed) rat being trapped is enough to elicit door-opening, we antagonized outward behavioral displays of distress with metyrapone (mty), which blocks autonomic signs of distress without altering somatomotor behavior. Rats so treated do not show distress but otherwise appear normal; these animals were not helped.

## Methods

All experimental procedures were reviewed and approved by the University of Chicago Institutional Animal Care and Use Committee (IACUC).

### Subjects

Two-month-old Sprague-Dawley (SD) rats were used for all experiments (total N=392). Rats used in the anesthesia experiments were purchased from Harlan (Indianapolis, IN, United States; N=44), and the others were purchased from Charles River (Portage, MI, United States, N=348). All rats were male and were housed in pairs. Rats had *ad libitum* access to chow and water in a 12:12 light-dark cycle and were given two weeks to acclimate to the housing environment and their cagemate.

### Habituation

Two weeks after arriving at the animal facility, animals were habituated to the testing rooms, experimenters (who were kept constant for each cohort of rats), and testing arenas. Testing arenas were constructed of acrylic (50 × 50 cm, 32-60 cm high) and were kept constant for each pair or rats. On day 1 of habituation, rats were transported by cart to the testing room and left undisturbed in their home cages on the cart. Thereafter, rats were transported to the room and left undisturbed for 15 min prior to further habituation procedures. On day 2, rats were briefly handled. Starting with the second day of habituation, rats were weighed 3 times weekly for the duration of the experiment; no animal lost weight during the experiment. On days 3-6, rats were handled for 5 minutes by each experimenter and then placed together (in housing pairs) in the testing arenas for 30 minutes. After each habituation session, rats were returned to their home cages and to the housing room. Within each cage, rats were randomly chosen to be either the free or trapped rat. Rats did not switch roles.

To habituate rats to intraperitoneal (i.p.) injections and to minimize stress related to the injection itself, rats to be injected in the testing phase received i.p. saline injections once daily for at least 5 days preceding testing. After receiving saline injections, rats were habituated to the testing arenas for 30 minutes as above. No free rat received habituating injections.

### Open field testing

On the day following completion of habituation, rats were placed individually in an arena for 30 min and their activity recorded (open field testing). The arenas were the same as were used during habituation, but open field testing represented the first time each rat was in the arena alone. Open field testing was used as a noninvasive metric of individual rat behavior.

As it turns out, no differences were seen and data from open field testing are not included in this report. Nonetheless, the animals experienced this testing, and we include it to provide a complete account of the rats’ treatment.

### Protocol for injection

To test the effect of mdz-treated trapped rats, injected rats were divided into four treatment groups (high dose mdz, 2.0 mg/kg; low dose mdz, 1.25 mg/kg; Saline; Uninjected; all i.p.). After the injections, rats sat in their home cages for 15 minutes for the mdz to take effect (rats in the Saline and Uninjected groups also sat for 15 minutes). Testing started immediately after this 15-minute interval. Note that the doses of mdz used lessen the affective display of distress in rats (e.g. Ben-Ami Bartal et al 2016; McGregor et al 2004; Cruz et al 1994) but are not sedating (Ben-Ami Bartal et al 2016).

Rats in the Anesthesia group were injected with Euthasol (Virbac Corporation, Fort Worth, TX, USA) diluted with deionized water to a final concentration of 39 mg/ml sodium pentobarbital and administered at a dose of 40 mg/kg, i.p.. After the injection, rats were actively monitored for vital signs (breathing and heart rate) and signs of general anesthesia (absence of righting reflex; absence of spontaneous movement). Injected rats that did not fully lose consciousness after 15 minutes were given another injection with 0.1 ml injection solution (3.9 mg sodium pentobarbital). Rats that were not anesthetized after the second injection were excluded from testing on that day. After all injected rats were unconscious or excluded, testing started.

Complete general anesthesia was successfully induced in 94 of 96 sessions. Two make-up sessions were then added at the end of the experiment to make up for the two failures. In four sessions, the trapped rat was dead at the end of testing and was subsequently replaced with a male Sprague Dawley rat of similar age and size. Finally, in a small minority of sessions (8/96, 8%) the trapped rat emerged from anesthesia during the experiment.

To further test the effect of sedation, a group of rats received a sedating dose of mdz (4.0 mg/kg, i.p.). These animals were also tested 15 minutes after the injection.

Rats assigned to the Metyrapone group were injected with mty (Cayman Chemical, Ann Arbor, Michigan, USA), administered at a dose of 25 mg/kg subcutaneously (s.c.). For the Metyrapone group, rats were left undisturbed for 30 minutes after injection for the drug to take effect. Animals in a control group received a saline injection s.c. and were also left undisturbed for 30 minutes—rats in this group are referred to as Saline-Metyrapone.

Mty blocks corticosterone synthesis in the adrenal cortex (Jenkins et al 1958). At the dose used (25 mg/kg s.c.), mty eliminates the increase in corticosterone associated with restraint stress (Calvo et al 1998) and reverses stress-induced physiological responses such as increases in body temperature and spontaneous locomotor activity (Drouet et al 2010). Yet, mty has no effect on basal plasma corticosterone in non-stressed rats (Roozendaal et al 1996; Calvo et al 1998).

### Trapped rat paradigm

On each testing day, rats were transported to the testing room and left undisturbed in their home cage for 15 minutes. Then rats were colored with markers to permit tracking individual rats’ movements. The free rat was colored red and the trapped rat colored blue. Rats were weighed and when indicated injected. After the appropriate waiting period, if any, the trapped rat was placed inside a restrainer which was positioned in the arena center. Restrainers are clear acrylic tubes (25 × 8.75 × 7.5 cm; Harvard Apparatus, Holliston, MA) with several slits, allowing for visual, olfactory, auditory, and vibrissal communication between rats. The free rat (the trapped rat’s cagemate) was then placed in the arena and allowed to roam freely. The door to the restrainer could only be opened from the outside and therefore only by the free rat. If the free rat did not open the restrainer door by 40 min, the investigator opened the restrainer door “halfway,” to a 45° angle, greatly facilitating door-opening by either rat. Only door-openings that occurred prior to the halfway opening were counted as such.

Rat dyads always remained in the arena for a full hour. Testing sessions were repeated once per day for 12 days. All sessions were run during the rats’ light cycle between 0800 and 1730. After each session, rats were returned to their home cages and the arena and restrainer were washed with 1% acetic acid followed by surface cleaner.

Some trapped rats (n=32, 38%) succeeded in opening the door from inside the restrainer. When this happened, the trapped rat was placed immediately back in the restrainer, and an acrylic blocker was inserted, preventing his access to the door. If the free rat subsequently opened the door, the blocker was removed, allowing the trapped rat to exit the restrainer. The blocker was then used for that trapped rat on all following test days. If the free rat failed to open the door by 40 min, the blocker was removed when the door was opened halfway.

### Reverse-door paradigm

Procedures in the reverse-door paradigm were the same as those in the trapped rat paradigm except that 1) the door was reversed so that it could only be opened from the inside; and 3) half of the rats were placed in the restrainer without a free rat outside and half were tested with a witness rat.

### Empty Restrainer Experiment

For the empty restrainer experiment, a free rat was put in the arena with an empty restrainer with its door facing the outside. As in the standard paradigm, the door was opened half-way at 40 minutes. One rat opened on the final day of planned testing. In order to determine whether that rat was reinforced by this experience, we tested that rat and the remaining group on the following (13^th^) day.

### Latency and statistical analysis

For each animal, a set of metrics were calculated including the number of openings (Openings), the number of consecutive openings (Consecutives), the day of the first opening (First-opening-day) and the total time until opening (Minutes-to-opening). For free rats that never opened over the 12 days of testing, the first day of opening was set to 13. A cutoff time of 40 minutes was assigned to rats that did not open on any given day. Analyses included all animals except where stated that analyses were restricted to rats that opened.

An ideally reinforced animal opens the restrainer on all subsequent opportunities after the first opening. As a measure of deviation from ideal reinforcement, we calculated the difference between the number of openings and the number of consecutive openings (Openings-minus-consecutives); the number and length (in days) of breaks (Breaks) was also noted. For additional metrics of learning and reinforcement, we subtracted the timing of the third opening from that of the first in terms of days (Third-minus-first-day) and latency (in minutes, Third-minus-first-latency). Animals that never opened (termed non-openers) were not included in reinforcement analyses.

All analyses were carried out in R.

All data are presented and illustrated as median and IQR (25th to 75th percentile). When appropriate, a Mann-Whitney U test was used to compare metrics between two groups, and a Kruskal-Wallis rank sum test was used between more than two groups. Post-hoc analyses were investigated through Dunn’s pairwise test with Bonferroni correction. The P values for all comparisons are stated.

### Principal component analysis

A principal component analysis was performed to identify patterns describing opening variability (Supplemental methods, table S1). We identified three principal components that describe most of the variability (>90%) between groups. These principal components are a linear combination of just a few variables. Because of the high proportion of variability accounted for by a small number of variables, raw variables were used in lieu of principal component values to aid interpretability. We used one raw metric to construct dissimilarity matrices for initiation (First-opening-day) and two for reinforcement dissimilarity matrices (Breaks and Third-minus-first-latency).

Dissimilarity matrices were used to determine both within- and between-group variability. In short, each animal is a point on a one-dimensional line (for the single initiation metric) or in two-dimensional space (for the two reinforcement metrics). Each value in the matrix represents the median Euclidean distance between animals in each pair of groups for between-group variability, or between animals in a single group for within-group variability (for detailed explanation see Supplemental methods). Euclidean distances were min-max normalized so that the greatest distance present in the entire dataset (for each variable) was set to one.

## Results

### Initiation and reinforcement are two separate processes in the helping paradigm

A principal component analysis on openings from all conditions identified three principal components, each of which accounts for more than 10% of the variability, and cumulatively account for more than 90% of the variability in the data (table S1). The first principal component identified the total number of openings (Openings) and the number of consecutive day openings (Consecutives). These metrics reflect overall performance (PC1).

PC2 and PC3 reflect the processes of initiation and reinforcement, respectively. The day of first opening (First-opening-day), and total minutes until opening (Minutes-to-opening; summed across the days) are metrics that reflect initiation (PC2). Since these two variables are highly correlated (r > 0.99), one metric - First-opening-day - was chosen as the metric for initiation hereafter. The number of breaks in opening (Breaks; an opening followed by no opening on the next day), and the difference between the third and first opening latencies (Third-minus-first-latency) are metrics that describe reinforcement (PC3). Thus, we use Breaks and Third-minus-first-latency, which are not correlated to each other (r < 0.01) as metrics of reinforcement.

The small number of metrics identified by principal component analysis were used to construct dissimilarity matrices that inform initiation and reinforcement. For each set of conditions, heat maps of dissimilarity matrices were made from selected metrics to analyze initiation and reinforcement; these matrices are shown in panels D and E, respectively, of Figures 1-3 and 5-6.

**Figure 1:**
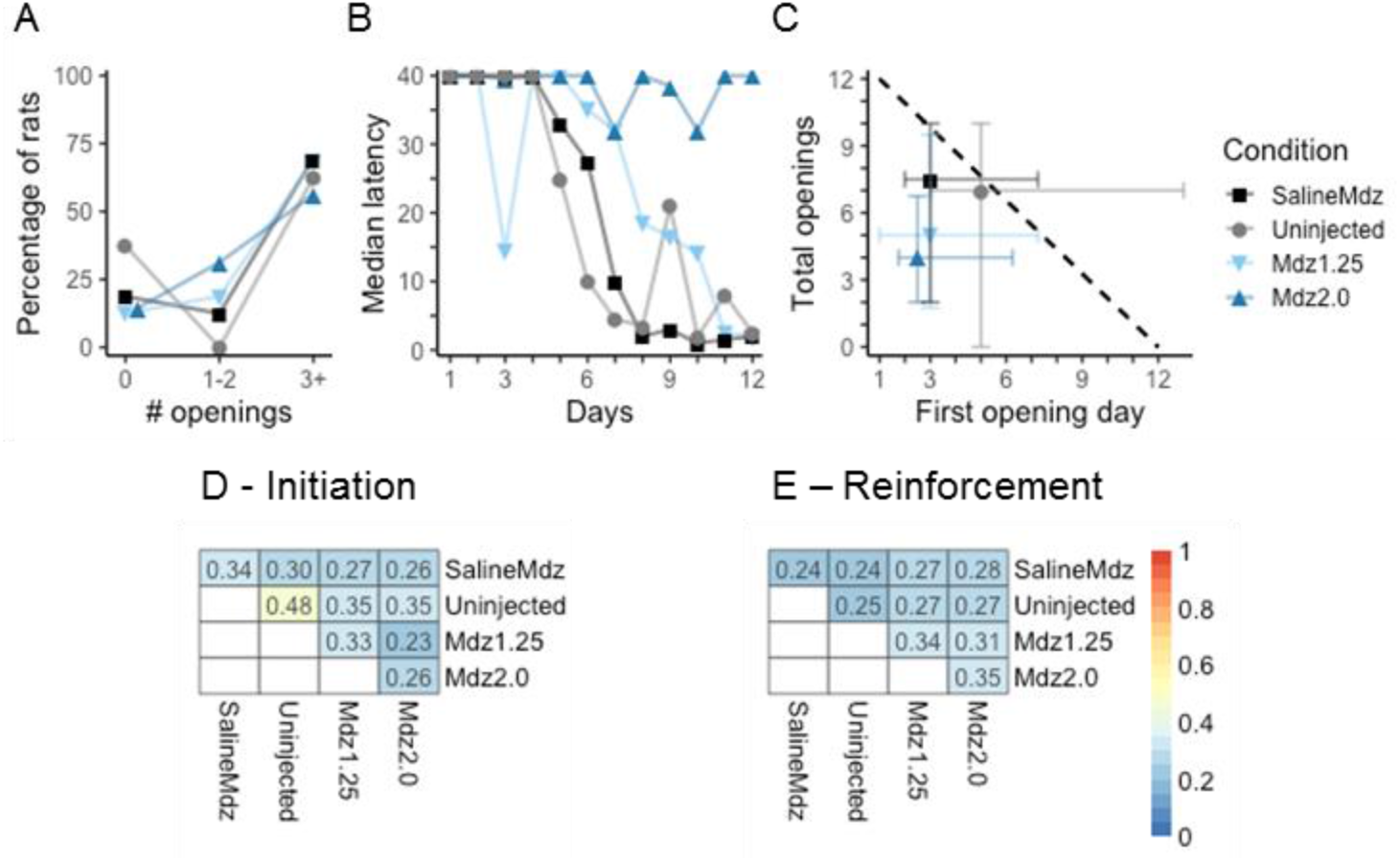
Free rats open only slightly more for control rats (Saline and Uninjected) than for mdz-treated animals. (A) Number of rats in each group that opened three times or more, one or two times, or never opened is shown. (B) Median latency for each group across the 12 experimental days. (C) The first opening day (x-axis), a metric of initiation is compared to the total openings (y-axis). The black dashed line denotes perfect reinforcement. Animals in control groups are nearly perfectly reinforced, whereas those treated with mdz are less so. (D-E) Dissimilarity matrices summarize within (diagonal cells) and between (off diagonal cells) group variability. (D) Initiation similarity scores do not vary considerably between control and mdz-treated groups. (E) With respect to reinforcement metrics, control animals were more similar to all groups than mdz-treated animals were to themselves.

One additional illustration of reinforcement is evident in Panel C which graphs First-opening-day against (total) Openings. For a perfectly reinforced animal, Openings will be equal to 13 minus First-opening-day, plotted as the dotted line. The horizontal distance from the condition’s median First-opening-day to the dotted line is another reflection of reinforcement, with values closest to the line representing the most reinforced conditions.

### Anxiolytic injection attenuates helping reinforcement

To test whether reducing a trapped rat’s affective distress reduces helping behavior, we administered mdz to trapped rats. Pairs of rats were divided into four groups (each N=16) with trapped rats that received high dose mdz (Mdz2.0), low dose mdz (Mdz1.25), saline (Saline), or no injection (Uninjected; see Methods for doses and procedural details). Each trapped rat was paired with its cagemate who served as the free rat.

Mdz-treatment had modest effects on door-opening (Fig. 1). When comparing all groups, no differences were observed in initiation (First-opening-day: χ^2^(3, N = 64) = 3.4, P = 0.34) or reinforcement (Openings-minus-consecutive: χ^2^(3, N = 51) = 1.5, P = 0.68; Third-minus-first-latency: χ^2^(3, N = 51) = 0.75, P = 0.86). However, upon pooling rats into Control-Mdz (Saline + Uninjected) and Mdz (Mdz1.25 + Mdz2.0) groups, it appears that rats injected with mdz opened for the first time slightly earlier than control animals (Control-Mdz: day 4.5, IQR 2-13; Mdz: day 3, IQR 1-7; Mann-Whitney U(N = 64) = 633, P = 0.05, r^2^ = 0.04). The slight advance in First-opening-day after mdz-treatment has been observed in rats accessing chocolate (Ben-Ami Bartal et al 2016), and is likely due to an anxiolytic release of the rat’s fear of venturing into the arena center.

### Anxiolytic injection in trapped rats does not block self-release in reverse-door paradigm

The asymmetric finding that mdz given to the free (Ben-Ami Bartal et al 2016), but not to the trapped rat (above), blocks helping may result from an asymmetry in the effects of mdz on free and trapped rats. A reasonable possibility is that due to the greater stress occasioned by being trapped in a tube compared to bearing witness to another’s restraint, a given dose of mdz will block affective arousal less completely in the trapped than the free rat. To determine the effect of mdz on the trapped rat’s distress, we placed trapped rats in a restrainer with a door that could only be opened from the inside (the reverse-door paradigm). This allowed us to indirectly measure the stress experienced by rats as the latency of self-release (an inverse metric) and the consistency of self-release across testing days (a direct metric).

Paralleling the first experiment, trapped rats in the reverse-door experiment received one of two doses of mdz (1.25 or 2.0 mg/kg), saline, or no injection. To compare the relative influences of being trapped and the potential for interaction with another rat, rats in each condition were tested either with a free rat present or not. Thus, this experiment had 4*2 levels: 4 treatment conditions (Reverse-Mdz2.0/ Reverse-Mdz1.25/ Reverse-Saline/ Reverse-Uninjected) and 2 free rat conditions (denoted by a suffix: “+” for with free rat, “-” for without free rat). Each level contained 16 trapped rats (N=128 total). Neither dose of mdz produced sedation (Ben-Ami Bartal et al 2016).

Most rats (119/128, 93.0%) opened the reverse-door and did so on most days (1023/1536 rat*opportunities; 66.6%). This homogeneity was reflected in the low median Euclidean distance between all pairs both within and between groups (Fig. 2D, E).

**Figure 2:**
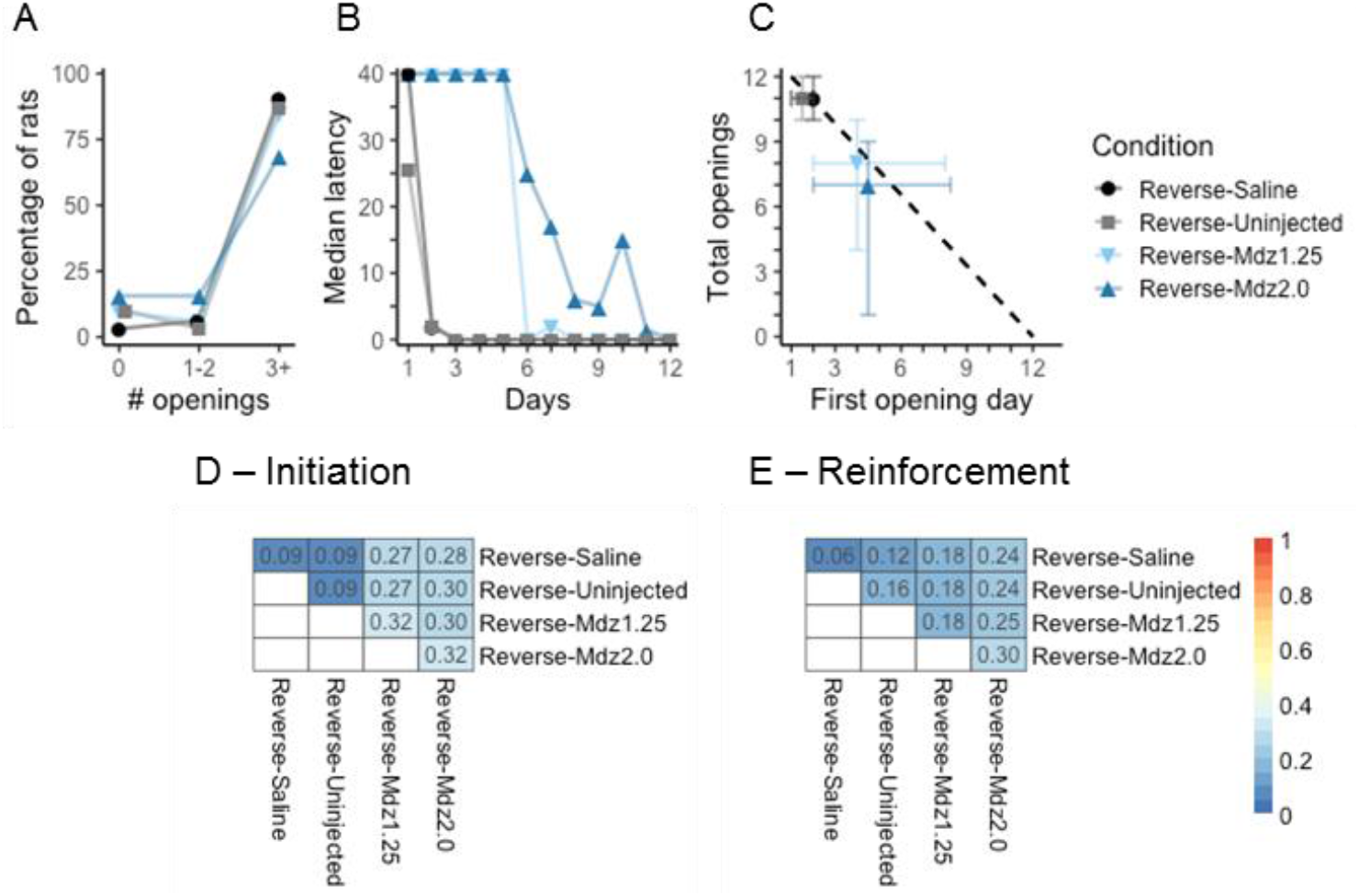
In the reverse-door paradigm, mdz-treated rats still released themselves from a restrainer at high rates. All panels have the same formats as in Figure 1. (A) Virtually all animals in all conditions released themselves at least three times. (B) Animals in control conditions (Reverse-Saline + Reverse-Uninjected) tend to initiate self-release earlier than those treated with mdz. (C) Similar to controls, animals treated with mdz who ever opened were nearly perfectly reinforced. (D – E) Groups were more similar in metrics of reinforcement than in metrics of initiation, with control groups more similar than mdz-treated groups.

Treatment affected the First-opening-day, a metric of initiation (Figure 2C; First-opening-day: χ^2^(7, N = 128) = 35.9, P < 0.001, η2 = 0; Dunn’s pairwise tests with Bonferroni corrections: Reverse-Saline+ vs Reverse-Mdz1.25- p = 0.001, Reverse-Saline+ vs Reverse-Mdz2.0-, p = 0.02, Reverse-Saline+ vs Reverse-Mdz2.0+ p = 0.005; Reverse-Uninjected-vs Reverse-Mdz1.25-, p = 0.008; Reverse-Uninjected+ vs Reverse-Mdz1.25-, p = 0.003; Reverse-Uninjected+ vs Reverse-Mdz2.0+, p = 0.01). Reverse-Saline and Reverse-Uninjected animals, grouped together hereafter as Reverse-Control, started to release themselves earlier (median 2, IQR 1-2) than did those treated with either dose of mdz (median 4, IQR 2-8; Mann-Whitney U(N = 128) = 892, P < 0.001, r^2^ = 0.25). The First-opening-day was similar in Reverse-Control animals regardless of whether a free rat was present. Of note, the presence of a free rat hastened the First-opening-day for rats receiving the lower dose of mdz (Reverse-Mdz1.25+: 2.5, 2-6.3; Reverse-Mdz1.25-: 6.5, 3-9; Mann-Whitney U(N = 32) = 178, P = 0.03,, r^2^ = 0.11).

Rats in all groups were highly reinforced following the first opening (Figure 2C, E). Thus, after opening once, rats opened on 907/1027 (88.3%) subsequent opportunities. Further, nearly all animals in all conditions consistently released themselves at very short latency (typically immediately, assigned 0 s) by the final days of testing (Figure 2B).

In sum, being trapped in a tube is a strong motivator for self-release among rats. This motivation is mildly reduced by mdz treatment and largely unaffected by the presence or absence of a witnessing rat (see table S2).

### An anesthetized rat elicits door-opening

Since the distress of trapped rats was never entirely blocked by low to moderate doses of mdz, we were unable to use mdz to determine whether helping requires some non-zero amount of trapped rat distress. To address this question by completely eliminating affective cues, we treated one group of trapped rats with a general anesthetic (Anesthesia) and a second group with a sedating dose of mdz (Mdz4.0; 4 mg/kg). Rats in both the Anesthesia and Mdz4.0 groups did not move and thus these groups are collectively termed the Immobile group. Opening for Immobile trapped rats was compared to the opening of an empty restrainer (Empty). Any difference observed can then be attributed to cognitive cues derived from the presence of the rat but not to any affective communication from that trapped rat. We also compared opening for Immobile rats to that observed in the Control-Mdz condition.

Free rats opened the door to an empty restrainer only once in 104 sessions (1%). In contrast, free rats opened for both Anesthesia and Mdz4.0 rats nearly as well as they opened for control trapped rats (Figure 3A-C). Pairwise differences confirmed that for opening initiation, the empty condition was radically different from both Control-Mdz and Immobile conditions which were not different from each other (First-opening-day: χ2(3, N = **56**) = 12.9, P = 0.005, η2 = 0.19, Dunn’s pairwise tests with Bonferroni corrections: Control-Mdz vs Empty, P = 0.003, Anesthesia vs Empty, P = 0.02, Mdz4.0 vs Empty, P = 0.005; Control-Mdz vs Anesthesia, P = 1.00; Control-Mdz vs Mdz4.0, P = 1.00; Anesthesia vs Mdz4.0, P = 1.00). There was no overall difference in reinforcement between Control-Mdz, Anesthesia and Mdz4.0 conditions (Openings minus Consecutive openings: Kruskal-Wallis rank sum χ2(3, N = 39) = 0.8, P = 0.86; Third latency minus first latency: χ2(3, N = 39) = 5.7, P = 0.12).

**Figure 3:**
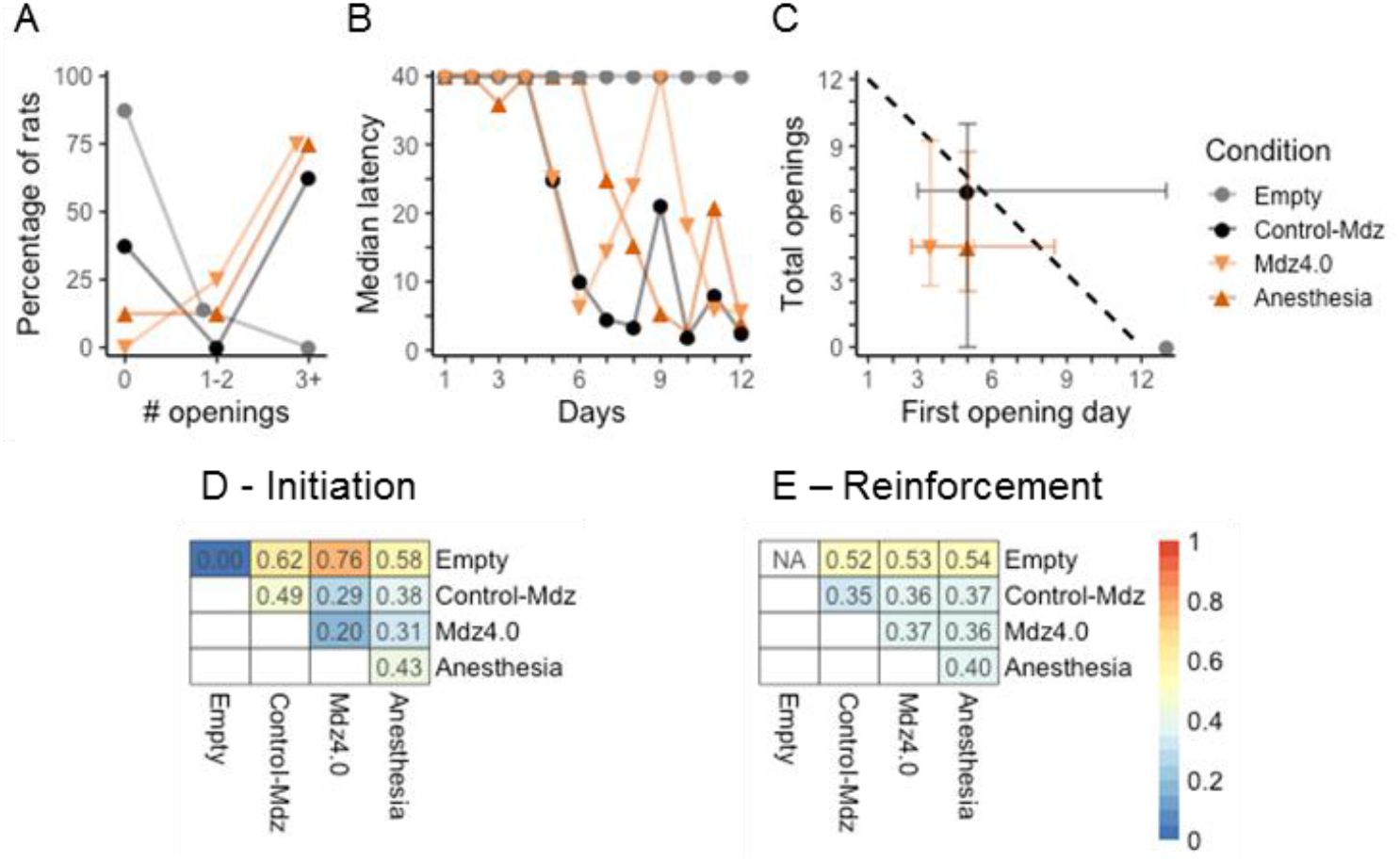
Cognitive cues from a trapped rat can elicit helping. All panels use the same formats as in Figure 1. (A-C) Free rats opened the door for anesthetized (Anesthesia) and heavily sedated (Mdz4.0) animals, but not for empty restrainers (Empty). Since only a single opening happened for the empty restrainer group, reinforcement was not considered. Anesthetized and heavily sedated animals were similar in terms of both initiation and reinforcement. (D) Rats opened for trapped Control-Mdz (Saline + Uninjected as in Fig. 1) and Immobile (Anesthesia + Mdz4.0) rats but not for an empty restrainer. (E) Rats opening for Control-Mdz and Immobile rats were reinforced in spite of high within and between group variability.

It should be noted that there was only one opening in the Empty condition and that opening occurred on the 12^th^ (intended to be final) day of testing. To determine if this was the start of a reinforced behavior, all rats in the Empty condition were tested for a 13^th^ day. No rat, including the rat that opened on the 12^th^ day, opened. Thus, there is no evidence that rats find opening for an empty restrainer reinforcing.

In sum, a rat, even one that is not communicating emotional cues, elicits door-opening whereas a restrainer does not.

### Rats exhibit specific behaviors towards Immobile conspecifics

To understand what drove opening for Immobile rats, we analyzed the behaviors exhibited by untreated rats toward Immobile rats and compared them to behaviors elicited by saline-injected and uninjected rats. After free rats opened the restrainer door, they exhibited a number of specific behaviors such as biting the testes or tail of the trapped rat and pulling the anesthetized rat out of the restrainer. The free rat then dragged the anesthetized rat out of the arena center with the ultimate effect of relocating the anesthetized rat off to the side, typically within a corner. These behaviors were other-oriented and appeared solicitous in nature.

Since dragging behavior has never occurred previously despite testing hundreds of pairs of rats in the trapped rat paradigm, we speculated that the behavior was elicited by the rats’ immobile state rather than by his trapped predicament. To test this idea, we placed anesthetized rats in the open (not in a restrainer) in the center of the arena. We compared the behavior of free rats toward unrestrained, anesthetized rats to the behavior of free rats toward unrestrained rats receiving an injection of either mdz or saline.

Accordingly, we studied four groups (each N=8) of Sprague-Dawley rats who received i.p. injections of saline, low-dose mdz (1.25 mg/kg), high-dose mdz (2.0 mg/kg), or of a general anesthetic. The injected rat was placed in the center of an open arena and an uninjected stranger was placed in the periphery of the arena. The pair was allowed to interact for 10 minutes. We manually coded for four solicitous behaviors (Fig. 4) that we observed in the last experiment.

**Figure 4.**
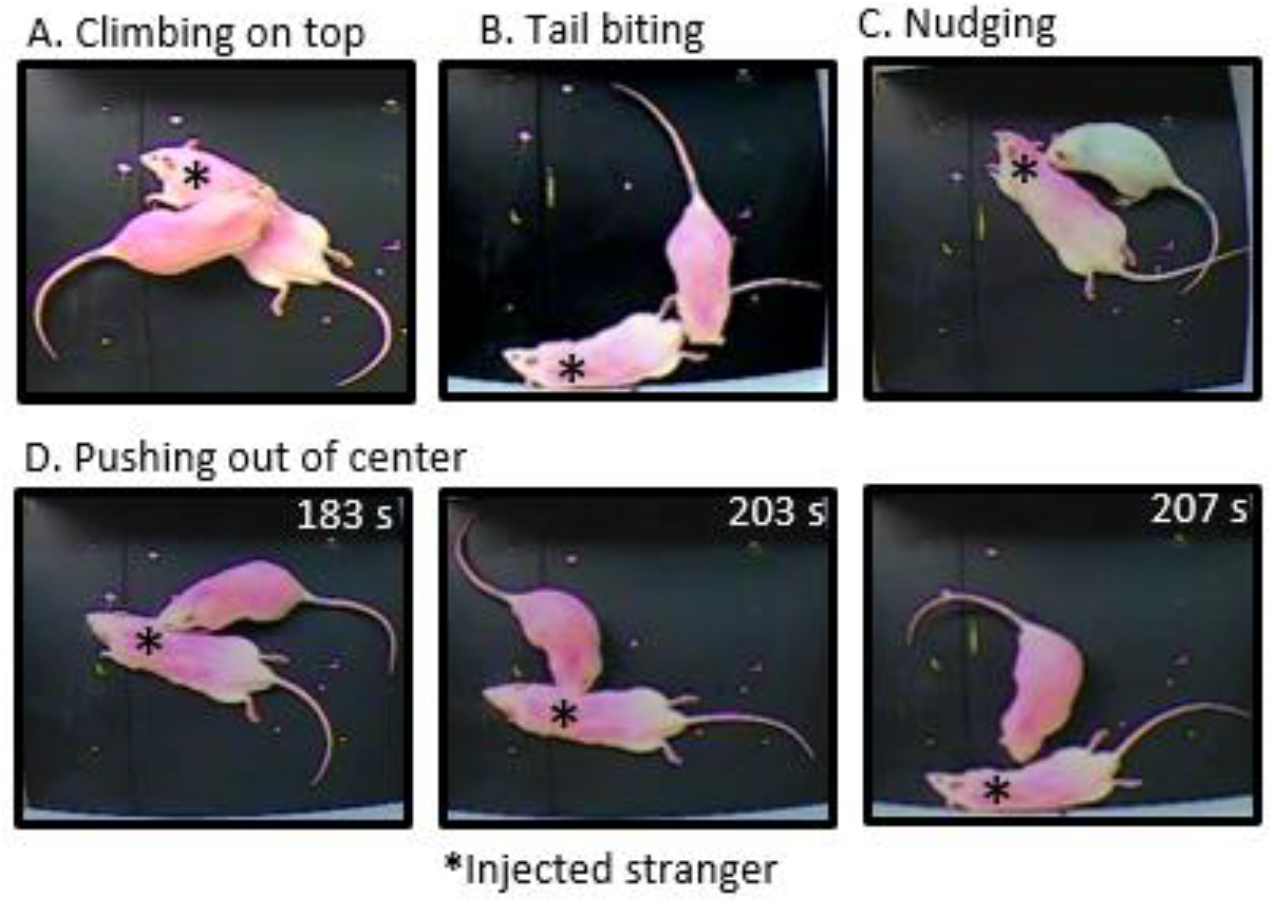
Immobilized rats (asterisks) elicit solicitous behaviors. Examples of climbing on top (A), tail biting (B), nudging (C) and a sequence of images (see time stamps in upper right) showing an immobilized rat being pushed out of the center.

The proportion of uninjected rats in each group that exhibited any of the concerned behaviors (see Fig. 4) was then calculated (see Table 1). Pushing rats to the arena wall, nudging, and tail biting occurred more frequently for anesthetized rats than for saline-treated rats (Fisher’s exact test: climbing p=0.28; pushing out of center p=0.03; nudging p=0.04; tail biting p=0.004). Pushing rats to the arena wall and tail-biting were also more frequently observed for anesthetized rats than for mdz-treated rats (Fisher’s exact test: pushing out of center p=0.03; tail biting: p=0.04).

**Table 1.**
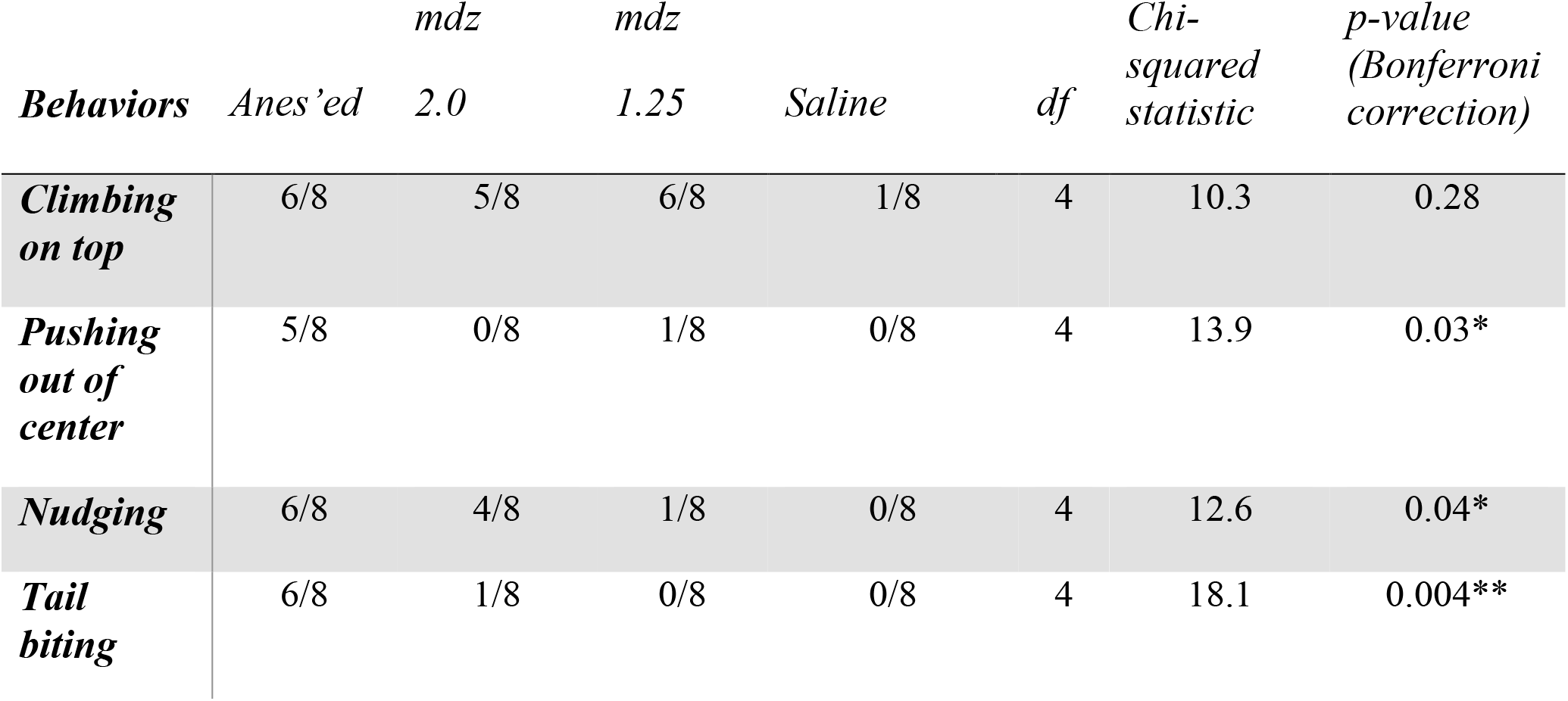
The proportions of rats exhibiting four specific behaviors toward rats treated as listed in column titles. The details of the chi-squared tests for each behavior are listed in the table, with the p-value corrected for multiple comparisons with the Bonferroni correction. Abbreviation: *df*: degrees of freedom.

### Diminished stress response from trapped rats, without cognitive cues suggestive of impairment, does not elicit helping behavior

Having observed that immobility elicited solicitous behavior even in the absence of a rat being trapped, we next asked whether the situational cue of a rat being trapped was sufficient to elicit door-opening. To isolate the trapped situation from other cognitive and affective cues, we used mty to selectively block corticosterone (CORT) synthesis in the trapped rat. Mty produces no conspicuous behavioral effects but blocks affective distress displays. To the human eye, mty-treated trapped rats (Metyrapone group) exhibited no observable distress signals but, unlike Immobile rats, moved normally. Mty was administered 30 min before the start of testing whereas mdz was administered 15 min prior; accordingly, we added a control group (N=8) that received saline 30 mins before testing (Saline-Metyrapone) rather than 15 min before as was the case in the Saline group above.

Overall, mty-treatment of trapped rats greatly reduced Openings from 72% (69/96) for the Saline-Metyrapone group to 30% (29/96). Only 2 out of 8 rats in the Metyrapone group (25%) opened the restrainer on consecutive days whereas 7 of 8 (87.5%) rats in the Saline-Metyrapone group did. Dissimilarity matrices showed that mty-treatment greatly altered reinforcement with less effect on initiation (Fig. 5D, E). Free rats consistently opened for trapped Saline-Metyrapone rats starting on day 3 (IQR 2-3.25) and thereafter were nearly perfectly reinforced (Fig. 5C). Thus, the Saline-Metyrapone group was similar to other control groups where rats were tested with untreated or saline-treated trapped rats. The median First-opening-day for the Metyrapone group was 3.5 (IQR 1.8-13) which was sandwiched between that of the Saline-Metyrapone (median 3, IQR 2-3.3; Mann-Whitney U(N = 16) = 35, P = 0.65, r^2^ = 0.004) and Control-Mdz (median 4, IQR 2-13) groups.

**Figure 5:**
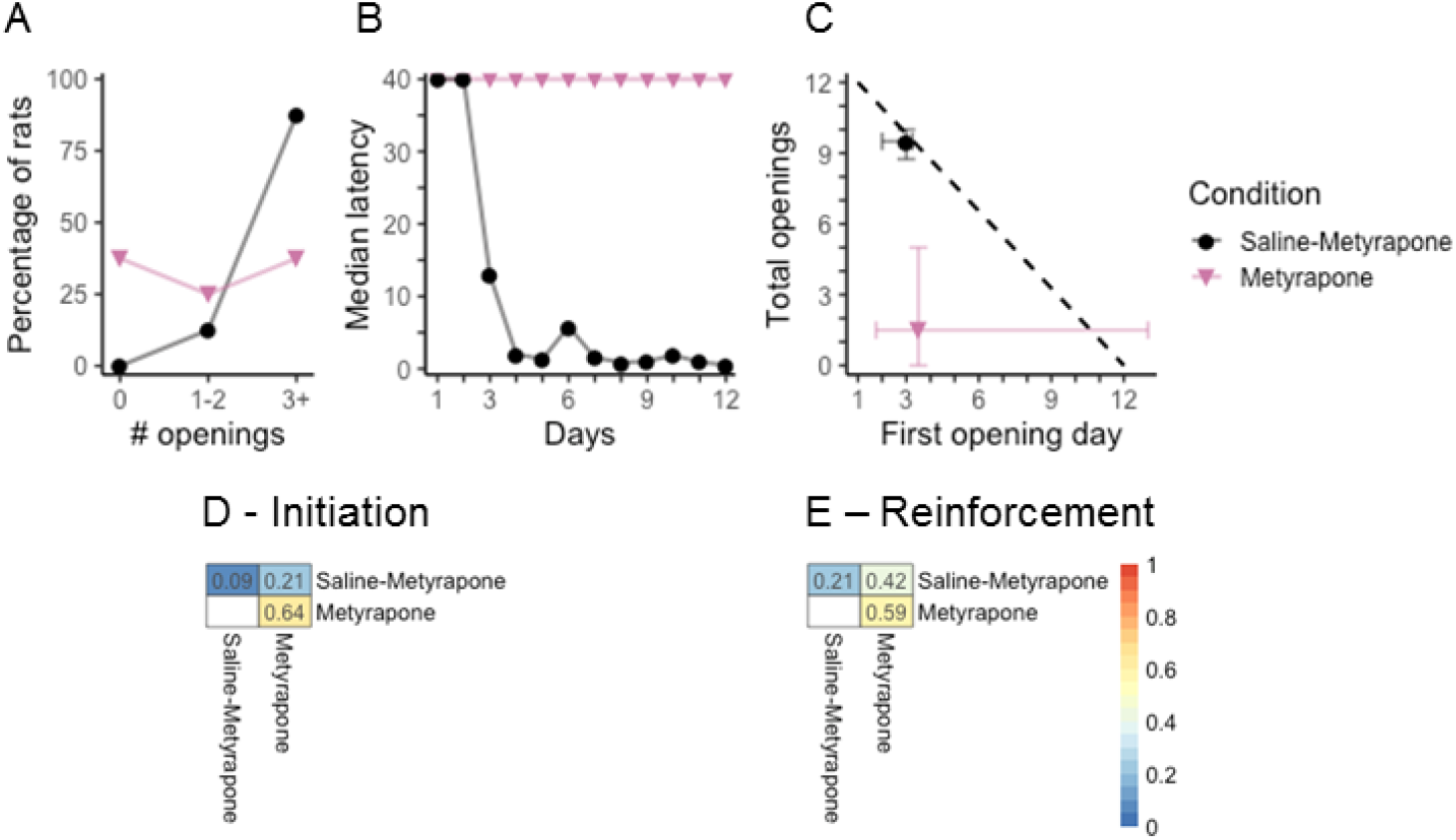
Control animals display more consistent helping than mty treated animals. All panels use the same formats as in Figure 1. (A-C) Rats consistently initiated opening for trapped Saline-Metyrapone rats and were nearly perfectly reinforced, whereas openings for trapped Metyrapone animals were rare and not reinforcing. (D-E) Initiation and reinforcement were highly variable for the Metyrapone condition.

It should be noted that free rats tested with mty-treated trapped rats behaved heterogeneously. Six of these animals never opened on consecutive days; the remaining two opened consistently (for 11 and 12 days). Thus, the median Euclidean distance for the Metyrapone group is near or above 0.6 for reinforcement and initiation, respectively (Fig. 5D, E).

### Comparison of all groups with a free rat

To make overall comparisons of all groups involving the potential for a free rat to open the restrainer door (this excludes the Reverse-door conditions), we made three super-groups to compare to the Metyrapone and Empty groups which each stood on their own. The Control-all group includes 40 rat pairs where the trapped animals did not receive a pharmacologically active agent (16 Uninjected, 16 Saline, 8 Saline-Metyrapone). The Immobile group consists of pairs with trapped animals that received either 4.0 mg/kg mdz or general anesthetic. Finally, pairs with trapped animals receiving either 1.25 or 2.0 mg/kg of mdz comprise the Mdz group as mentioned above.

Comparing across these five groups, we found that the empty group, which was uniform in behavior (no within group variation), differed from all other groups. Simply put, rats did not open a restrainer door in the absence of a trapped rat in whatever physiological state.

Rats tested with Mdz or Immobile trapped rats showed only subtle differences from the Control-all group in initiation, suggesting that animals were motivated to start helping other rats even when affective cues were reduced or completely absent (Fig. 6B, D). Differences in reinforcement were similarly modest (Fig. 6C, E). In stark contrast, rats tested with Metyrapone trapped rats, displayed large within-group variability, and also differed the most from the other non-empty conditions. Differences were in both initiation and reinforcement, suggesting that the motivation to help is decreased when there are neither obvious behavioral cues (immobility) nor affective cues (outward distress).

**Figure 6:**
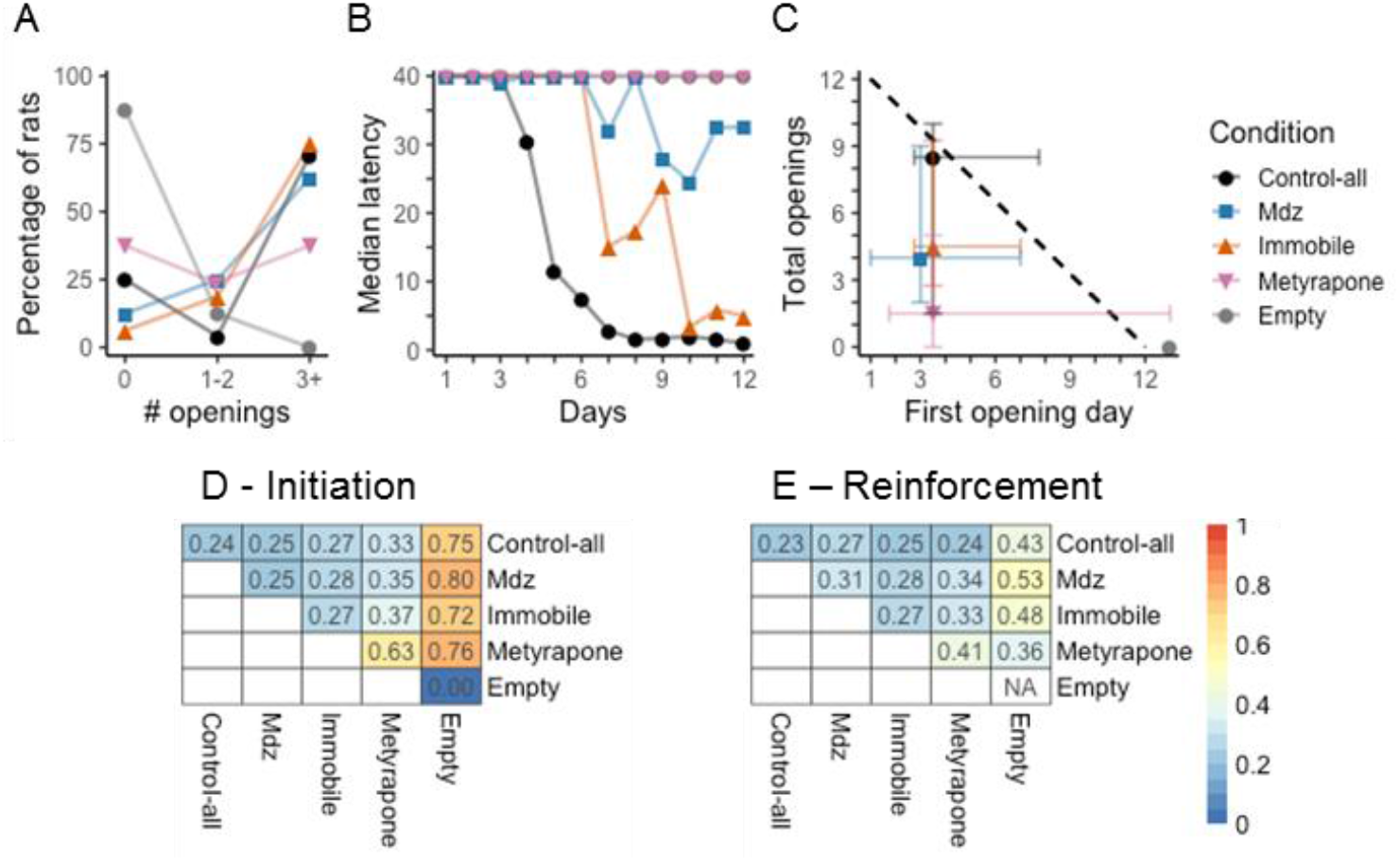
Control-all, immobilized, and mdz-treated animals behaved similarly. All panels use the same formats as in Figure 1. (A-C) Opening for Control-all animals (16 Uninjected + 16 Saline + 8 Saline-Metyrapone) is conspicuously different from opening for an empty restrainer. Openings for mdz-treated and for Immobile animals stand in an intermediate position between the extremes. Opening for mty-treated animals is more similar to that of rats tested with an empty restrainer than to any other group. (D) Dissimilarity scores demonstrate that initiation of opening for Control-all, Immobile and Mdz rats are nearly uniform. Interestingly, within-group variability is lower than between-group variability for the Metyrapone condition. This is also reflected in the IQR shown in panel C for the Metyrapone group. (E) All conditions differed from the Empty condition in reinforcement. Note that one animal in the Empty condition opened and is compared to animals in the other conditions and yet because he was the only one, no intra-group metric was calculated. Reinforcement for Metyrapone trapped rats was highly variable with a lack of reinforcement for 3/5 rats that were ever opened for.

## Discussion

### Current findings

Here we show that either affective or cognitive cues are sufficient to elicit helping in rats (Table 2). Neither is necessary in that either one suffices in the absence of the other. We further demonstrate that not all cognitive cues elicit helping in rats. An anesthetized rat elicits helping but the situation of being trapped, in the absence of frank emotional cues, rarely does. Thus, an immobile rat behaves at odds with the viewer’s life-long expectations. This motivates the viewer into other-oriented solicitous behavior including helping. We propose that concern arising from cognitive dissonance is a rudimentary form of cognitive empathy akin to emotional contagion being a rudimentary form of affective empathy. Yet, rats do not appear to use situational cues to motivate helping. Thus, while a rudimentary form of cognitive empathy obtains in rats, we find no evidence for more advanced forms of cognitive empathy, such as perspective-taking, in rats.

**Table 2.**
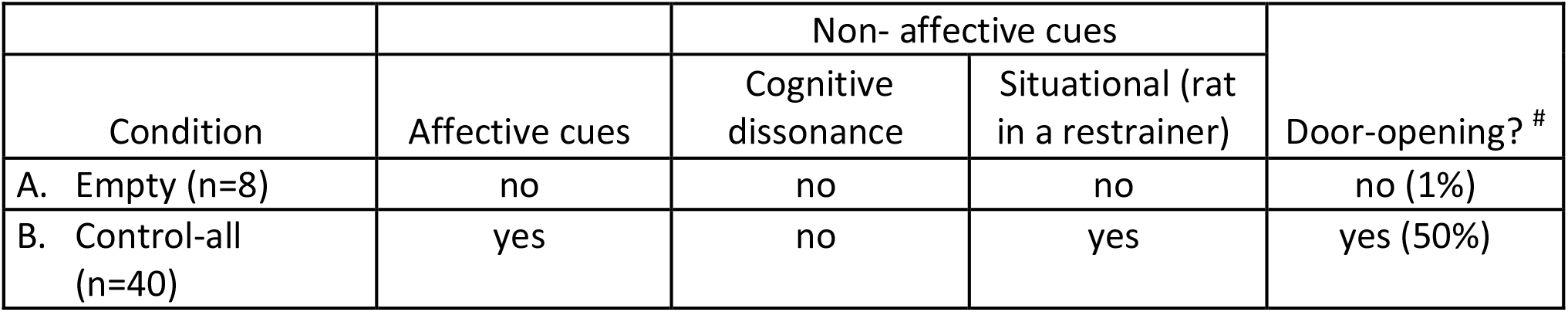
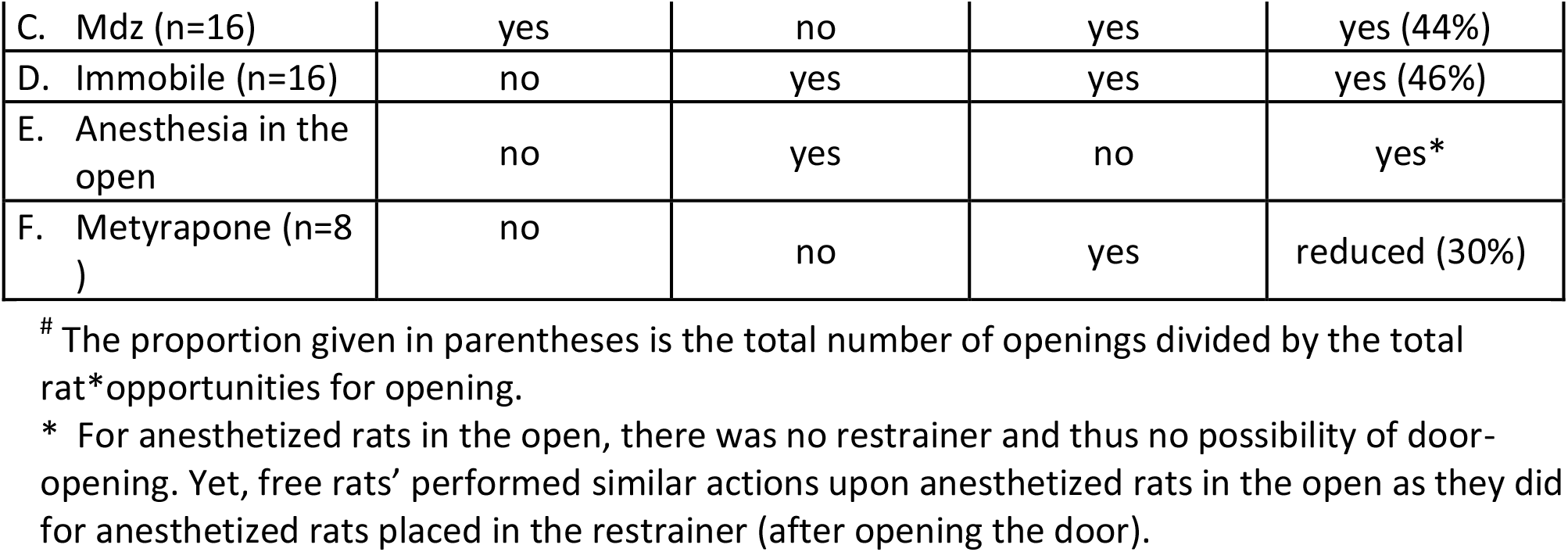
By comparing the outcomes resulting from specific combinations of affective, cognitive and situational cues available to the free rat in each condition (excluding reverse-door), several conclusions can be deduced. The conditions and comparisons leading to each conclusion are given in the parentheses with the letters referring to the conditions labeled in the table. Conclusions: 1) a restrainer is neither sufficient nor necessary to elicit door-opening (A); 2) either affective (B-F; C-F) or cognitively dissonant (D-F; E) cues are sufficient to elicit helping; 3) neither affective (D-F; E) nor cognitively dissonant cues (B-F; C-F) is necessary – either one suffices in the absence of the other; 4) situational cues are neither sufficient (F) nor necessary (E) to elicit helping or other similarly solicitous behaviors.

| Condition | Affective cues | Non- affective cues |  | Door-opening? <sup>#</sup> |
| --- | --- | --- | --- | --- |
|  |  | Cognitive dissonance | Situational (rat in a restrainer) |  |
| A. Empty (n=8) | no | no | no | no (1%) |
| B. Control-all (n=40) | yes | no | yes | yes (50%) |
| C. Mdz (n=16) | yes | no | yes | yes (44%) |
| D. Immobile (n=16) | no | yes | yes | yes (46%) |
| E. Anesthesia in the open | no | yes | no | yes* |
| F. Metyrapone (n=8 ) | no | no | yes | reduced (30%) |

### Being trapped is aversive

When rats were trapped with a reverse-door, they consistently released themselves from a closed restrainer and did so immediately (latency of 0 s) within a day or two of first opening. This direct evidence that rats would prefer not to be trapped in a restrainer is consistent with decades of work demonstrating restraint tubes as an aversive stressor (Buynitsky and Mostofsky 2009; Glavin et al 1994). Of note, the motivation for self-release was only altered by the presence of a witness rat when trapped rats were treated with mdz (table S2).

Rats treated with an anxiolytic dose of mdz began opening the reversed door later than did control rats. However, once they began, mdz-treated rats continued to open the door consistently and at short latency. Thus, anxiolytic treatment only mildly reduced the distress of trapped rats, primarily by delaying the onset of self-release. That mdz was largely ineffectual at blocking the distress of trapped rats informs interpretation of our finding that free rats open the door, at near control levels, for trapped rats treated with mdz. The minimal effect of anxiolytic treatment on the distress of trapped rats parallels a minimal effect on helping for anxiolytic-treated trapped rats, which is what we observed. These results are consistent with helper rats’ being motivated by the distress of trapped rats, suggestive of helping motivated by empathic concern.

### Free rats continue to open for mdz-treated rats

We previously showed that blunting affective processes in the free rat obliterates helping (Ben-Ami Bartal et al 2016). The doses used here were identical and the experiments were conducted concurrently. Thus, it would appear that the distress of the free rat is substantially less than that of the trapped rat. This is consistent with emotional contagion in which the free rat catches only some portion of the trapped rat’s total distress.

In light of trapped rats remaining distressed even after mdz treatment, the modest effects of mdz administered to the trapped rat on the free rat’s door-opening behavior are understandable. The trapped rat’s distress is so great that anxiolytic treatment reduces it insufficiently to substantially alter the free rat’s caught distress; as a result, door-opening still occurs. Thus, in the presence of affective distress, emotional contagion is the vehicle that leads to motivated helping.

The distress communicated through emotional contagion appears to summate across available sources. A recent study showed that rats integrate the emotional contagion derived directly from the trapped rat with information gained indirectly from bystanders’ reactions to the trapped rat (Havlik et al 2020). When the bystanders show concern, distress from emotional contagion adds up and helping occurs earlier and more often. Conversely, when bystanders are uninterested in the trapped rat, the lack of distress exhibited subtracts from the direct emotional contagion and helping occurs infrequently.

In sum, a trapped rat’s distress elicits emotional contagion that motivates a free rat to open the restrainer door. This is helping motivated by affective empathic concern.

### Cognitive cues can motivate solicitous interactions

Rats opened for anesthetized and heavily sedated animals (collectively termed Immobile) as they did for control animals, compelling evidence that opening can be motivated by something other than the trapped rat’s observable distress. Similarly, mice chew through a blocker to release a mouse trapped in a tube, regardless of whether the trapped mouse is anesthetized or not (Ueno et al 2019). Here, we found that even in the absence of a restrainer, rats nudged, bit, and otherwise interacted with immobile rats. While the proper semantic characterization for these behaviors - concern, consolation, caring - is debatable, its other-oriented nature is indisputable. Here we term the behavior as *solicitous* because it appears both other-oriented and positive, reflective of protectiveness rather than aggression.

Since being trapped was unnecessary to elicit solicitous behavior, other-oriented behaviors are a reaction to the immobile rat’s apparent state rather than to the situation. Immobile rats are remarkable for lacking both affect and posture, an appearance that would be unprecedented in a rat’s life. The witnessing rat’s door-opening and solicitous behavior clearly demonstrate that he views the immobile rat as arousing and motivating. The witness rat does ***not*** base his behavior upon affective contagion or motor mimicry.

The Perception-Action Model of Preston and de Waal (2002) holds that state-matching is required for empathic concern and ultimately targeted helping. Our results are in variance with this idea as solicitous and helping behavior occurred without state-matching in rats. Empathic actions occur without state-matching in humans as well. For example, one does not need to feel afraid to console a child who is afraid of an unfamiliar dog so that it is possible to care even in the absence of empathy (Bloom, p. 64).

How does a witnessing rat move from detection of the immobile rat as remarkable to motivated solicitous behavior? One possibility is that the witnessing rat is simply curious. In this formulation, the rat would perform solicitous behaviors without an affective motivation. This could be thought of as a social form of exploration. Alternatively, it may be that the witnessing rat translates the non-affective cues of the immobile rat into an aversive affective state that motivates other-oriented care. According to this model, the witnessing rat would translate the immobile rat’s cues into an affective state that shares none of the qualities displayed by the immobile rat. Instead, it would be an affect produced *de novo* in the witnessing rat. Such a transformation from non-affective cues to an affective motivation would constitute cognitive rather than affective empathy and would not be based upon emotional contagion.

Cognitively-motivated prosocial behavior has been reported before. Rats share food with other rats who are not food-deprived and thus not frankly hungry, a form of pro-social behavior that does not depend on a display of affect (Hernandez-Lallement, 2015; Márquez et al 2015). While food-sharing is initially motivated by cognitive cues, it may be sustained by a positive affective reaction as the sharing rat sees the other rat receive and eat the food. In contrast, immobile rats do not demonstrate any behavior in response to the solicitations of the active rat. Immobile rats do not even exit the restrainer unless pulled out by the active rat. In ligx ht of the lack of any feedback available from immobile rats, the remarkable consistency of the active rat’s solicitous behavior suggests internal reinforcement. Thus, the solicitous rat appears to gain an internal affective reward from his behavior. In effect, the solicitous rat may act to reduce a negative affect engendered by the immobile rat’s extraordinary appearance. These arguments favor the idea that the other-oriented behaviors exhibited toward immobile rats are a form of cognitive empathy rather than social curiosity.

### Mty serves to pharmacologically remove restrainer-initiated distress

Rats treated with mty, which blocks autonomic signs of distress by blocking corticosterone synthesis, showed neither distress nor odd behavior. Free (untreated) rats rarely opened for trapped rats treated with mty. It is as though mty pharmacologically removes the restrainer. Combined with our finding that untrapped saline-injected rats do not elicit concern, we conclude that rats are motivated by behavior but not by restraint *per se*.

Most mty-treated animals failed to elicit opening whereas 2/8 were consistently opened for. This heterogeneity is most likely related to a heterogeneity in drug efficacy such that the treatment did not completely block corticosterone synthesis in all animals. Of note, we used the smallest reported dose that suppresses stress-induced corticosterone synthesis, 25 mg/kg s.c., to avoid behavioral side effects such as changes in locomotion. (Roozendall et al., 1996; Calvo et al., 1998).

### Concern for immobile rats arises from cognitive dissonance

The entire trapping context was unnecessary to elicit solicitous behavior. Rats climbed on top of not-trapped immobile rats, pushed them around, often into a corner, and bit their tail. These behaviors were not elicited by untreated, saline-injected or even rats who received an affect-blunting dose of mdz, evidence that these behaviors constitute an other-oriented concern specifically elicited by the odd appearance and behavior of a motionless rat.

Other-oriented actions toward anesthetized rats appear motivated by the gross departure of an immobile rat’s behavior from expectations, arguably a rodent form of *cognitive dissonance*. This cognitive dissonance is born of the failure of an immobile rat to meet life-long predictions for conspecific appearance and behavior. A rat without affect that also lacks normal posture and shows no purposeful movement contradicts all of a rat’s past experience and thus his expectations. Nonetheless the motionless rat elicits opening. The equally unprecedented appearance of an empty restrainer does not elicit behavior, highlighting the salience of social surprises over inanimate ones.

While concern for an immobile rat appears to be a rudimentary form of cognitive empathy, it does not share features with advanced forms of cognitive empathy such as perspective-taking. Rats were less motivated to open the door for mty-treated rats, rats who behaved normally (no cognitive dissonance) and showed no affective distress (no emotional contagion). This demonstrates that free rats are not translating the situation of being trapped into a motivation for concern. Thus, rats do not show evidence of more advanced forms of cognitive empathy such as knowing another’s internal state or thoughts (for example from information shared through language), projecting oneself into another’s situation (“aesthetic empathy” used by writers for example), or imagining how one would feel if in another’s situation (“changing places in fancy”) (Batson 2009).

### Relative importance of affective, non-affective and situational cues

In sum, either affective or non-affective cues are sufficient to elicit other-oriented concern while neither is necessary. Situational cues alone were neither necessary nor sufficient to motivate prosocial behavior. Thus, rats may be motivated alternatively by affect or by cognitive dissonance, the latter comprising a rudimentary form of cognitive empathy.

## Supporting information

VieiraSuganoTrappedStatistics

VieiraSuganoTrappedRawData

## Acknowledgements

The assistance of M.S. Bernardez Sarria, W. Kwateng, M. Sales, M. Barajas, R. Zhou, is gratefully acknowledged. The authors thanks Inbal Ben-Ami Bartal for insightful comments on manuscript drafts. The authors declare no competing financial interests.

## References

Atsak, P, Orre M, Bakker P, Cerliani L, Roozendaal B, Gazzola V, Moita M, Keysers C (2011) Experience Modulates Vicarious Freezing in Rats: A Model for Empathy Ferrari PF, ed. PLoS ONE 6:e21855.

Ben-Ami Bartal I, Decety J, Mason P (2011) Empathy and Pro-Social Behavior in Rats. Science 334:1427–1430.

Ben-Ami Bartal I, Rodgers DA, Bernardez Sarria MS, Decety J, Mason P (2014) Pro-social behavior in rats is modulated by social experience. eLife 3:e01385.

Ben-Ami Bartal I, Shan H, Molasky NMR, Murray TM, Williams JZ, Decety J, Mason P (2016) Anxiolytic Treatment Impairs Helping Behavior in Rats. Front Psychol 7 Available at: http://journal.frontiersin.org/Article/10.3389/fpsyg.2016.00850/abstract [Accessed January 11, 2022].

Blair RJR (2005) Responding to the emotions of others: Dissociating forms of empathy through the study of typical and psychiatric populations. Consciousness and Cognition 14:698–718.

Bloom P (2016) Against empathy: the case for rational compassion, First edition. New York, NY: Ecco, an imprint of HarperCollins Publishers 48.

Burkett JP, Andari E, Johnson ZV, Curry DC, de Waal FBM, Young LJ (2016) Oxytocin-dependent consolation behavior in rodents. Science 351:375–378.

Buynitsky T, Mostofsky DI (2009) Restraint stress in biobehavioral research: Recent developments. Neuroscience & Biobehavioral Reviews 33:1089–1098.

Calvo N, Martijena ID, Molina VA, Volosin M (1998) Metyrapone pretreatment prevents the behavioral and neurochemical sequelae induced by stress. Brain Research 800:227–235.

Cox SS, Reichel CM (2020) Rats display empathic behavior independent of the opportunity for social interaction. Neuropsychopharmacol 45:1097–1104.

Cruz APM, Frei F, Graeff FG (1994) Ethopharmacological analysis of rat behavior on the elevated plus-maze. Pharmacology Biochemistry and Behavior 49:171–176.

Drouet J-B, Michel V, Peinnequin A, Alonso A, Fidier N, Maury R, Buguet A, Cespuglio R, Canini F (2010) Metyrapone blunts stress-induced hyperthermia and increased locomotor activity independently of glucocorticoids and neurosteroids. Psychoneuroendocrinology 35:1299–1310.

Dziobek I, Rogers K, Fleck S, Bahnemann M, Heekeren HR, Wolf OT, Convit A (2008) Dissociation of Cognitive and Emotional Empathy in Adults with Asperger Syndrome Using the Multifaceted Empathy Test (MET). J Autism Dev Disord 38:464–473.

Eslinger PJ (1998) Neurological and Neuropsychological Bases of Empathy. Eur Neurol 39:193–199.

Glavin GB, Paré WP, Sandbak T, Bakke H-K, Murison R (1994) Restraint stress in biomedical research: An update. Neuroscience & Biobehavioral Reviews 18:223–249.

Harari H, Shamay-Tsoory SG, Ravid M, Levkovitz Y (2010) Double dissociation between cognitive and affective empathy in borderline personality disorder. Psychiatry Research 175:277–279.

Havlik JL, Vieira Sugano YY, Jacobi MC, Kukreja RR, Jacobi JHC, Mason P (2020) The bystander effect in rats. Sci Adv 6:eabb4205.

Hernandez-Lallement J, van Wingerden M, Marx C, Srejic M, Kalenscher T (2015) Rats prefer mutual rewards in a prosocial choice task. Front Neurosci 8 Available at: http://journal.frontiersin.org/article/10.3389/fnins.2014.00443/abstract [Accessed January 11, 2022].

Jenkins JS, Meakin JW, Nelson DH, Thorn GW (1958) Inhibition of adrenal steroid 11-oxygenation in the dog. Science 128:478–480.

Jeon D, Kim S, Chetana M, Jo D, Ruley HE, Lin S-Y, Rabah D, Kinet J-P, Shin H-S (2010) Observational fear learning involves affective pain system and Cav1.2 Ca2+ channels in ACC. Nat Neurosci 13:482–488.

Langford DJ, Crager SE, Shehzad Z, Smith SB, Sotocinal SG, Levenstadt JS, Chanda ML, Levitin DJ, Mogil JS (2006) Social Modulation of Pain as Evidence for Empathy in Mice. Science 312:1967–1970.

Márquez C, Rennie SM, Costa DF, Moita MA (2015) Prosocial Choice in Rats Depends on Food-Seeking Behavior Displayed by Recipients. Current Biology 25:1736–1745.

McGregor IS (2004) Neural Correlates of Cat Odor-Induced Anxiety in Rats: Region-Specific Effects of the Benzodiazepine Midazolam. Journal of Neuroscience 24:4134–4144.

Meyza KZ, Bartal IB-A, Monfils MH, Panksepp JB, Knapska E (2017) The roots of empathy: Through the lens of rodent models. Neuroscience & Biobehavioral Reviews 76:216–234.

Panksepp JB, Lahvis GP (2011) Rodent empathy and affective neuroscience. Neuroscience & Biobehavioral Reviews 35:1864–1875.

Preston SD, de Waal FBM (2002) Empathy: Its ultimate and proximate bases. Behav Brain Sci 25:1–20.

Roozendaall B, Bohus B, McGaugh JL (1996) Dose-dependent suppression of adrenocortical activity with metyrapone: Effects on emotion and memory. Psychoneuroendocrinology 21:681–693.

Sato N, Tan L, Tate K, Okada M (2015) Rats demonstrate helping behavior toward a soaked conspecific. Anim Cogn 18:1039–1047.

Ueno H, Suemitsu S, Murakami S, Kitamura N, Wani K, Matsumoto Y, Okamoto M, Ishihara T (2019) Helping-Like Behaviour in Mice Towards Conspecifics Constrained Inside Tubes. Sci Rep 9:5817.

