## Supplemental Materials for "Helping can be driven by non-affective cues in rat"

**Raw data**

Of the 392 rats used, 232 were tested subjects; the remainder were trapped rats. Data collected from the subject rats are available in the accompanying excel file TrappedAllData.xlsx. Each subject (row) is a rat, and each condition is a specific treatment. Data consists of latency of door opening for all 12 testing days. The cut-off latency of 40 minutes indicate that an animal did not open during that testing session. All other metrics are calculated derivatives of these numbers.

**Data summary**

Summary statistics for each condition, as well as larger groups made from different conditions (e.g., Control-all, Immobile) are present in the accompanying excel file Statistics.xlsx. Description of groups as well as calculated metrics are described in the file.

Animals that opened only once and never opened again were not reinforced at all. However, the score in the Openings-minus-consecutive metric (1 opening minus 0 consecutive openings) would be one for these animals, suggestive of perfect reinforcement. For that reason, a score of 5 was assigned to animals that opened only once and never opened again. 5 is the highest score observed for an animal with multiple openings and bad reinforcement, and thus is a conservative lower estimation for an animal that was not reinforced at all.

**Data normalization**

For principal component analysis and dissimilarity matrices, all variables were normalized by min-max scaling $(X_{n}-X_{min})/ (X_{max}-X_{min})$, so that their range is 0-1 while their distribution is unchanged. Normalization was done across all the rats (N = 232)

**Principal component analysis**

A principal component analysis using the *prcomp* function (R version 1.4.1717) across animals in all conditions identified three principal components, each of which accounts for more than 10% of the variability, and cumulatively account for more than 90% of the variability in the data (Table 1). The first principal component identified the total number of openings (Openings) and the number of consecutive day openings (Consecutives). These metrics reflect overall performance (PC1).

PC2 and PC3 reflect the processes of initiation and reinforcement, respectively. The day of first opening (First-opening-day), and total minutes until opening (Minutes-to-opening; summed across the days) are metrics that reflect initiation (PC2). Since these two variables are highly correlated (r > 0.99), First-opening-day is used as a metric of initiation hereafter. The number of breaks in opening (Breaks; an opening followed by no opening on the next day), and difference between the third and first opening latencies (Third-minus-first-latency) are metrics that describe reinforcement (PC3). Opening-minus-consecutives and Breaks are highly correlated (r = 0.90); Thus, we use Breaks and Third-minus-first-latency, which are not correlated to each other (r < 0.01) as metrics of reinforcement.

| **Variable** | **PC1** | **PC2** | **PC3** | **PC4** | **PC5** | **PC6** | **PC7** | **PC8** | **PC9** | **PC10** | **PC11** |
| --- | --- | --- | --- | --- | --- | --- | --- | --- | --- | --- | --- |
| Openings | **-0.43** | -0.16 | -0.07 | 0.13 | -0.02 | -0.04 | -0.34 | 0.14 | -0.09 | -0.72 | 0.32 |
| First opening day | 0.13 | **0.67** | -0.03 | 0.14 | 0.03 | -0.02 | -0.12 | -0.03 | 0.58 | -0.04 | 0.40 |
| Consecutive | **-0.49** | -0.10 | 0.15 | 0.06 | -0.05 | -0.12 | -0.17 | 0.30 | -0.04 | 0.64 | 0.42 |
| Breaks | 0.25 | -0.18 | **-0.51** | 0.15 | 0.16 | -0.03 | 0.19 | 0.72 | 0.16 | 0.00 | 0.00 |
| Total breaks | 0.36 | -0.21 | 0.09 | -0.21 | 0.00 | 0.06 | 0.40 | -0.13 | -0.23 | -0.08 | 0.74 |
| Openings - consecutive | 0.17 | -0.15 | **-0.61** | 0.19 | 0.08 | 0.21 | -0.46 | -0.42 | -0.14 | 0.23 | 0.15 |
| Third - first Day | 0.29 | -0.13 | 0.29 | -0.18 | -0.38 | 0.61 | -0.41 | 0.30 | 0.12 | 0.00 | 0.00 |
| Third - first Latency | -0.12 | 0.05 | **-0.42** | -0.56 | -0.64 | -0.27 | 0.04 | -0.03 | 0.12 | 0.00 | 0.00 |
| Second - first Latency | -0.23 | 0.10 | -0.11 | -0.69 | 0.61 | 0.28 | -0.07 | 0.04 | 0.05 | 0.00 | 0.00 |
| Last 6 Openings | -0.42 | 0.15 | -0.21 | 0.22 | -0.21 | 0.65 | 0.49 | -0.07 | -0.02 | 0.00 | 0.00 |
| Minutes to Opening | 0.12 | **0.60** | -0.07 | -0.03 | -0.07 | -0.01 | -0.12 | 0.27 | -0.73 | 0.00 | 0.00 |
| **Cumulative Proportion** | 58% | 81% | 92% | 97% | 98% | 100% | 100% | 100% | 100% | 100% | 100% |

table S1. Rotation matrix summarizing the principal component analysis. PC1, PC2 and PC3 account for more than 90% of the variability on the data and are highly correlated with metrics of overall performance (Openings, Consecutive), metrics of initiation (First opening day, Minutes-to-opening) and metrics of reinforcement (Breaks, Openings-minus-consecutive, Third-minus-first-Latency).

|  | **Openings all 12 days** | **Rats that never opened** | **Openings after first opening** |
| --- | --- | --- | --- |
| Reverse-Mdz1.25 | 56% | 9% | 86% |
| Reverse-Mdz2.0 | 48% | 16% | 77% |
| Reverse-Saline | 83% | 3% | 95% |
| Reverse-Uninjected | 79% | 9% | 91% |
| Control + | 73% | 13% | 86% |
| Control - | 79% | 6% | 94% |
| MDZ + | 44% | 19% | 75% |
| MDZ - | 60% | 6% | 87% |

table S2. Opening performance is compared across the four reverse conditions.

**Dissimilarity matrices**

Dissimilarity matrices were created to obtain a numerical estimator for within and between group variability. Using the variables describes above (Minutes-to-opening for initiation, Breaks and Third-minus-first-latency for reinforcement), each animal can be represented in a one or two dimensional space. Then, Euclidean distance between every pair of animals was calculated using the in R. Then, median distances were used to summarize group distances. Within-group distance was calculated as the median distance between pairs of animals from the same condition, whereas between-group distance was calculated as the median distance between pairs of animals from different conditions. Lastly, the median distance was normalized such that both initiation and reinforcement distances range between 0 (least distant, all animals are the same) and 1 (most distant).

Euclidean distances were calculated between every pair of animals. The median distance between all animals within the same group was used as a measure of in-group similarity. Similarly, the median distance between all pairs of animals from two groups measured between-group similarity. All data are presented in heatmaps created with the pheatmap package (Kolde, 2019).

**References**

Raivo Kolde (2019). pheatmap: Pretty Heatmaps. R package version 1.0.12. CRAN - Package pheatmap
